# Maternal exposure to the cannabinoid agonist WIN 55,12,2 during lactation induces lasting behavioral and synaptic alterations in the rat adult offspring of both sexes

**DOI:** 10.1101/2020.04.09.025932

**Authors:** Andrew F Scheyer, Milene Borsoi, Anne-Laure Pelissier-Alicot, Olivier JJ Manzoni

## Abstract

Consumption of cannabis during pregnancy and the lactation period is a rising public health concern (Scheyer et al., 2019). We have previously shown that exposure to synthetic or plant-derived cannabinoids via lactation disrupts perinatal programming of the gamma-aminobutyric acid trajectory in the prefrontal cortex and early-life behaviors (Scheyer et al., 2020b). Recently, we described lasting behavioral and neuronal consequences of Δ9-tetrahydrocannabinol (THC) perinatal exposure via lactation (Scheyer et al., 2020a).

Here, we extend upon these findings by testing the effects in offspring of maternal exposure to the synthetic cannabinoid agonist WIN 55,12,2 (WIN). The data demonstrate that rats exposed during lactation to WIN display social, cognitive and motivational deficits at adulthood. These behavioral changes were paralleled by a specific loss of endocannabinoid-mediated long-term depression in the prefrontal cortex and nucleus accumbens, while other forms of synaptic plasticity remained intact. Thus, similarly to THC, perinatal WIN exposure via lactation induces behavioral and synaptic abnormalities lasting into adulthood.

## Introduction

Cannabis consumption by pregnant women is progressively increasing 1–4. The principle psychoactive component of cannabis, Δ9-tetrahydrocannabinol (THC), in addition to other cannabinoids, is actively transferred to the developing infant via breastfeeding (Hurd et al., 2019; Scheyer et al., 2019, 2020a, 2020b). During the perinatal period (i.e. during prenatal and early postnatal development), the developing brain is acutely sensitive to exogenous cannabinoids (Scheyer et al., 2019). We have previously shown that exposure to cannabinoids via lactation alters the developmental trajectory of the prefrontal cortex (PFC), which has been identified as a cortical hub essential to planning, cognitive flexibility and emotional behaviors (Goldman-Rakic, 1990) and a common target in various endocannabinoid-related synaptopathies (Araque et al., 2017). Our work revealed that exposure of lactating females to either THC, or a synthetic agonist of CB1R, altered the maturational trajectory of GABAergic transmission and led to behavioral abnormalities in early life (Scheyer et al., 2020b). Very recently, we reported that perinatal THC exposure via lactation elicited lasting, deleterious impacts on social behavior and synaptic plasticity in the PFC of the adult offspring (Scheyer et al., 2020a).

Here, we extend upon these findings by investigating the effects in both sexes of adult offspring of maternal exposure to the synthetic cannabinoid agonist, WIN 55,12,2 (WIN). These data demonstrate that rats exposed during lactation to WIN display social, cognitive and motivational deficits at adulthood. These behavioral changes were paralleled by a specific loss of endocannabinoid-mediated long-term depression in the PFC and nucleus accumbens, while other forms of synaptic plasticity remained intact. Thus, similarly to THC, perinatal WIN exposure via lactation induces behavior and synaptic abnormalities lasting into adulthood.

## Materials and Methods

### Animals

Animals were treated in compliance with the European Communities Council Directive (86/609/EEC) and the United States NIH Guide for the Care and Use of Laboratory Animals. The French Ethical committee authorized the project “Exposition Périnatale aux cannabimimétiques” (APAFIS #18476-2019022510121076 v3). All rats were group-housed with 12h light/dark cycles with ad libitum access to food and water. All behavioral, biochemical and synaptic plasticity experiments were performed on male and female RjHan:wi-Wistar rats (>P90) from pregnant females obtained from Janvier Labs. Pregnant dams arrived at E15 and remained undisturbed until delivery. Newborn litters found before 05:00p.m. were considered to be born that day (P0). Dams were injected daily subcutaneously (s.c.) from P01-10 with WIN (0.5 mg/kg/day), dissolved in 10% polyethylene glycol/10% Tween/80% saline and injected subcutaneously as described previously (Borsoi et al., 2019)(Scheyer et al., 2020b). Control dams (Sham) received vehicle.

### Behavioral procedures

#### Open field

Observations were conducted after rats were adapted to the room laboratory conditions for at least 1 h prior to testing. Tests were conducted in a 45 × 45 cm transparent Plexiglass arena. All behavioral procedures were performed between 10:00 am and 3:00 pm. A video tracking system (Ethovision XT, Noldus Information Technology) recorded the total distance traveled and time spent in the central zone (21 × 21 cm) of the apparatus (Borsoi et al., 2019; (Scheyer et al., 2020b).

#### Social interaction

the procedure was performed as previously (Scheyer et al., 2020b). The apparatus consisted of a transparent acrylic chamber (120 × 80 cm) divided into three equal compartments (40 cm each) partially separated by white walls. The central compartment was empty and lateral compartments had an empty wire cage (20 cm diameter) were an object or a new rat (social stimulus) were placed during the test. WIN or sham-exposed rats were individually habituated to the test cage containing the two empty wire cages for 5 min immediately prior to testing. The first trial (social approach, 5 min duration) consisted of giving the tested rat the option to socialize with either a novel object or a new, naïve, age- and sex-mate conspecific rat that were placed into the wire cages positioned on the arena’s opposite sides. Thirty minutes later, the tested rat returned to the apparatus for the second trial (social memory, 5min duration) wherein the two compartments held either the now-familiar rat from the first testing phase or a second, previously unknown, naïve, age- and sex-mate conspecific. Only rats with no compartment preference during the habituation phase were used. Time spent in each compartment and time spent exploring wire cages during the social approach and social memory phases were scored. Social Preference Ratio was calculated as time spent exploring either the wire cage containing the object, or the new rat divided by total time exploring both wire cages. Likewise, Social Memory Ratio was calculated as time spent exploring either the wire cage containing the rat used in the first trial or the new rat divided by total time exploring both wire cages. Recognition index higher than 0.5 indicates preferable object recognition memory.

#### Novel object recognition

the procedure was performed as previously (Borsoi et al., 2019). The test was performed in the same apparatus used in the open field test. It comprised training (acquisition trial) and test (5min duration each) phases. During the acquisition trial, the rat was placed into the arena containing two identical sample objects (A1 and A2) placed near the two corners at either end of one side of the arena (8 cm from each adjacent wall). Thirty minutes later, the rat returned to the apparatus containing two objects, one of them was a copy to the object used in the acquisition trial (A3), and the other one was novel (B). The objects in the test were placed in the same positions as during the acquisition trial. The positions of the objects in the test and the objects used as novel or familiar were counterbalanced between the animals. Exploration was scored when the animal was observed sniffing or touching the object with the nose and/or forepaws. Sitting on objects was not considered to indicate exploratory behavior. The recognition index was calculated as follow: time spent by each rat exploring the novel object divided by the total time spent exploring both objects. Recognition index higher than 0.5 indicates preferable object recognition memory.

#### Anhedonia

We performed sucrose consumption tests (Monleon et al., 1995; Bessa et al., 2009). Rats were exposed for 24 h to a bottle containing a sucrose solution (5% in tap water, Sigma), placed in the wire-top cage cover adjacent to standard tap water, followed by 12 h of water deprivation and a 20 minute exposure to two identical bottles (one filled with 5% sucrose solution and the other with water). Bottles were placed at opposite ends of the cage and counterbalanced across groups to avoid side bias. Sucrose preference was calculated as the ratio of the volume of sucrose versus volume or consumed during the 20-minute test. All animals were habituated to the testing room 24 h prior to initiating the sucrose preference test.

### Slice preparation

Adult male and female rats were anesthetized with isoflurane and sacrificed as previously described (Bara et al., 2018; Borsoi et al., 2019). The brain was sliced (300 µm) in the coronal plane with a vibratome (Integraslice, Campden Instruments) in a sucrose-based solution at 4°C (in mm as follows: 87 NaCl, 75 sucrose, 25 glucose, 2.5 KCl, 4 MgCl_2_, 0.5 CaCl_2_, 23 NaHCO_3_ and 1.25 NaH_2_PO_4_). Immediately after cutting, slices containing the medial prefrontal cortex (PFC) or the nucleus accumbens (NAc) were stored for 1 hr at 32°C in a low-calcium ACSF that contained (in mm) as follows: 130 NaCl, 11 glucose, 2.5 KCl, 2.4 MgCl_2_, 1.2 CaCl_2_, 23 NaHCO_3_, 1.2 NaH2PO_4_, and were equilibrated with 95% O2/5% CO_2_ and then at room temperature until the time of recording. During the recording, slices were placed in the recording chamber and superfused at 2 ml/min with low Ca^2+^ or normal Ca^2+^ ACSF (PFC and NAc respectively). All experiments were done at 32°C (PFC) or 25°C (NAc). The superfusion medium contained picrotoxin (100 mM) to block gamma-aminobutyric acid types A (GABA-A) receptors. All drugs were added at the final concentration to the superfusion medium.

### Electrophysiology

Whole cell patch-clamp of visualized layer five pyramidal medial PFC or medium spiny neurons and field potential recordings were made in coronal slices as previously described (Bara et al., 2018; Borsoi et al., 2019). Neurons were visualized using an upright microscope with infrared illumination. The intracellular solution was based on K+ gluconate (in mM: 145 K^+^ gluconate, 3 NaCl, 1 MgCl_2_, 1 EGTA, 0.3 CaCl_2_, 2 Na^2+^ ATP, and 0.3 Na^+^ GTP, 0.2 cAMP, buffered with 10 HEPES). The pH was adjusted to 7.2 and osmolarity to 290–300 mOsm. Electrode resistance was 4–6 MOhms. Recordings were performed with an Axopatch-200B amplifier as previously described (Bara et al., 2018; Borsoi et al., 2019)(Scheyer et al., 2020b). Data were low pass filtered at 2kHz, digitized (10 kHz, DigiData 1440A, Axon Instrument), collected using Clampex 10.2 and analyzed using Clampfit 10.2 (all from Molecular Device, Sunnyvale, USA).

A -2 mV hyperpolarizing pulse was applied before each evoked EPSC in order to evaluate the access resistance and those experiments in which this parameter changed >25% were rejected. Access resistance compensation was not used, and acceptable access resistance was <30 MOhms. The potential reference of the amplifier was adjusted to zero prior to breaking into the cell. Cells were held at -75mV.

Current-voltage (I-V) curves were made by a series of hyperpolarizing to depolarizing current steps immediately after breaking into the cell. Membrane resistance was estimated from the I–V curve around resting membrane potential (Martin et al., 2015). Field potential recordings were made in coronal slices containing the PFC or the NAc as previously described (Kasanetz et al., 2013). During the recording, slices were placed in the recording chamber and superfused at 2 ml/min with low Ca^2+^ ACSF. All experiments were done at 32°C. The superfusion medium contained picrotoxin (100 mM) to block GABA Type A (GABA-A) receptors. All drugs were added at the final concentration to the superfusion medium. The glutamatergic nature of the field EPSP (fEPSP) was systematically confirmed at the end of the experiments using the ionotropic glutamate receptor antagonist CNQX (20 mM), which specifically blocked the synaptic component without altering the non-synaptic.

Both fEPSP area and amplitude were analyzed. Stimulation was performed with a glass electrode filled with ACSF and the stimulus intensity was adjusted ∼60% of maximal intensity after performing an input–output curve (baseline EPSC amplitudes ranged between 50 and 150 pA). Stimulation frequency was set at 0.1 Hz.

### Data acquisition and analysis

The magnitude of plasticity was calculated at 0-10min and 30–40 min after induction (for TBS-LTP and eCB-LTD) or drug application (mGlu2/3-LTD) as percentage of baseline responses (Scheyer et al., 2020a). Statistical analysis of data was performed with Prism (GraphPad Software) using tests indicated in the main text after outlier subtraction (Grubb’s test, alpha level 0.05). All values are given as mean ±SEM, and statistical significance was set at p<0.05.

## Results

In rodent models, exposure to cannabinoids (both synthetic and plant-derived) during gestation or early development induces an array of deleterious consequences on behavior manifesting both at early life and adulthood (Hurd et al., 2019; Scheyer et al., 2019).

Previously, we have demonstrated that perinatal exposure via lactation to either the plant-derived phytocannabinoid Δ9-tetrahydrocannabinol (THC), or the synthetic cannabinoid, WIN 55,212-2 (WIN), induces a significant delay in the trajectory of GABAergic development in the PFC of developing offspring, an effect which is accompanied by substantial behavioral alterations (Scheyer et al., 2020b). Further, we recently reported that the progeny of dams similarly exposed via lactation to THC during the first 10 days of postnatal life exhibit lasting deficits in synaptic plasticity in the PFC as well as augmented social behavior at adulthood (Scheyer et al., 2020a).

Here, we used this same protocol of perinatal cannabinoid exposure in order to determine if synaptic and behavioral consequences are similarly produced following maternal administration of WIN. Thus, lactating dams were treated with a low dose of WIN (0.5 mg/kg, s.c.) or its vehicle (herein referred to as Sham) from postnatal day 1 to 10 (PND 1-10). Experiments were then conducted in the male and female offspring at adulthood (>PND90).

All treatment effects (e.g. Sham vs WIN) were found to be consistent across sexes. Thus, for figures and statistical analyses, data for male and female rats within treatment condition were combined. However, differences between the sexes within treatment conditions were noted in some measures. Details of within-treatment sex differences can be found in tables 1-9.

**Table 1.** Social approach and social memory data by sex.

| Condition | Test – Measure (unit) | Value | T-test (M v F) |
| --- | --- | --- | --- |
| Sham male | Social Preference (ratio) | 0.8647 ± 0.01743<br>N=15 | P = 0.5089 |
| Sham female | Social Preference (ratio) | 0.8375 ± 0.03569<br>N=8 |  |
| WIN male | Social Preference (ratio) | 0.9491 ± 0.01282<br>N=11 | P = 0.0255 |
| WIN female | Social Preference (ratio) | 0.8618 ± 0.03219<br>N=11 |  |
| Sham male | Social Memory (ratio) | 0.6147 ± 0.03886<br>N=15 | P = 0.0876 |
| Sham female | Social Memory (ratio) | 0.7463 ± 0.05970<br>N=8 |  |
| WIN male | Social Memory (ratio) | 0.6282 ± 0.05435<br>N=11 | P > 0.9999 |
| WIN female | Social Memory (ratio) | 0.6282 ± 0.03590<br>N=11 |  |
| Sham male | Social Preference – Time exploring object (sec) | 14.20 ± 1.853<br>N=15 | P = 0.8121 |
| Sham female | Social Preference – Time exploring object (sec) | 13.61 ± 1.610<br>N=8 |  |
| WIN male | Social Preference – Time exploring object (sec) | 6.586 ± 1.840<br>N=11 | P = 0.0420 |
| WIN female | Social Preference – Time exploring object (sec) | 16.05 ± 3.822<br>N=11 |  |
| Sham male | Social Preference – Time exploring rat (sec) | 95.38 ± 8.016<br>N=15 | P = 0.3569 |
| Sham female | Social Preference – Time exploring rat (sec) | 83.02 ± 10.25<br>N=8 |  |
| WIN male | Social Preference – Time exploring rat (sec) | 110.2 ± 9.669<br>N=11 | P = 0.4518 |
| WIN female | Social Preference – Time exploring rat (sec) | 100.8 ± 7.422<br>N=11 |  |
| Sham male | Social Memory – Time exploring familiar rat (sec) | 29.31 ± 3.733<br>N=15 | P = 0.2200 |
| Sham female | Social Memory – Time exploring familiar rat (sec) | 20.90 ± 5.375<br>N=8 |  |
| WIN male | Social Memory – Time exploring familiar rat (sec) | 26.17 ± 3.754<br>N=11 | P = 0.3621 |
| WIN female | Social Memory – Time exploring familiar rat (sec) | 35.52 ± 2.748<br>N=11 |  |
| Sham male | Social Memory – Time exploring novel rat (sec) | 47.06 ± 4.528<br>N=15 | P = 0.1116 |
| Sham female | Social Memory – Time exploring novel rat (sec) | 58.58 ± 5.159<br>N=8 |  |
| WIN male | Social Memory – Time exploring novel rat (sec) | 49.36 ± 8.196<br>N=11 | P = 0.7014 |
| WIN female | Social Memory – Time exploring novel rat (sec) | 53.04 ± 4.657<br>N=11 |  |

**Table 2.**
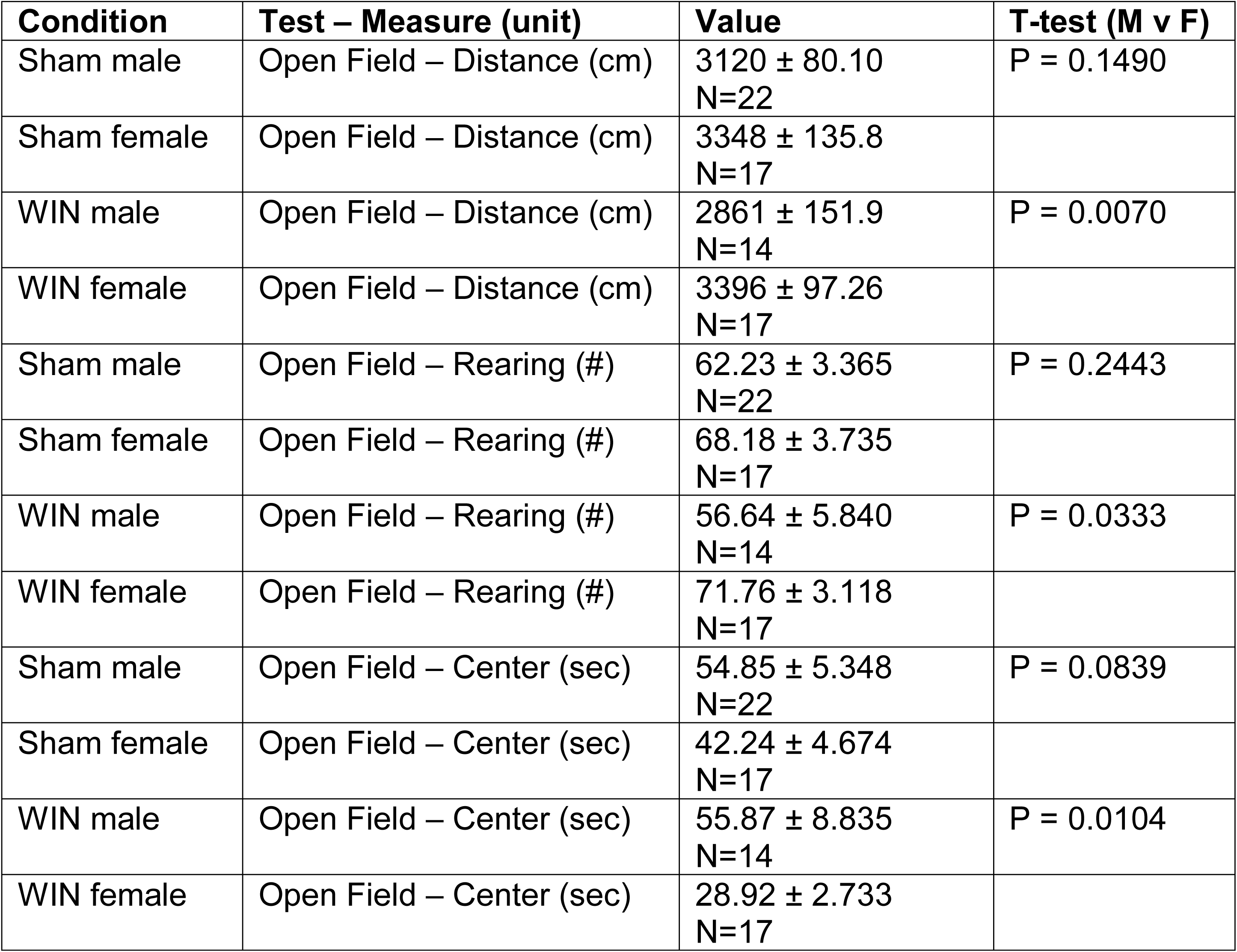
Open field data by sex.

**Table 3.** Novel Object Recognition data by sex.

| **Condition** | **Test – Measure (unit)** | **Value** | **T-test (M v F)** |
| --- | --- | --- | --- |
| Sham male | NOR test – Time spent on novel object (sec) | 12.75 ± 2.937<br>N=14 | P = 0.3813 |
| Sham female | NOR test – Time spent on novel object (sec) | 9.760 ± 1.582<br>N=8 |  |
| WIN male | NOR test – Time spent on novel object (sec) | 10.12 ± 1.972<br>N=9 | P = 0.2149 |
| WIN female | NOR – Time spent on novel object (sec) | 13.52 ± 1.777<br>N=17 |  |
| Sham male | NOR – Time spent on familiar object (sec) | 20.69 ± 2.940<br>N=14 | P = 0.1785 |
| Sham female | NOR – Time spent on familiar object (sec) | 14.60 ± 3.199<br>N=8 |  |
| WIN male | NOR – Time spent on familiar object (sec) | 11.67 ± 1.799<br>N=9 | P = 0.0101 |
| WIN female | NOR – Time spent on familiar object (sec) | 19.74 ± 2.258<br>N=17 |  |
| Sham male | NOR – Discrimination index | 0.6343 ± 0.02711<br>N=14 | P = 0.3287 |
| Sham female | NOR – Discrimination index | 0.5775 ± 0.04861<br>N=8 |  |
| WIN male | NOR – Discrimination index | 0.5550 ± 0.04213<br>N=8 | P = 0.5680 |
| WIN female | NOR – Discrimination index | 0.5888 ± 0.04015<br>N=17 |  |

**Table 4.**
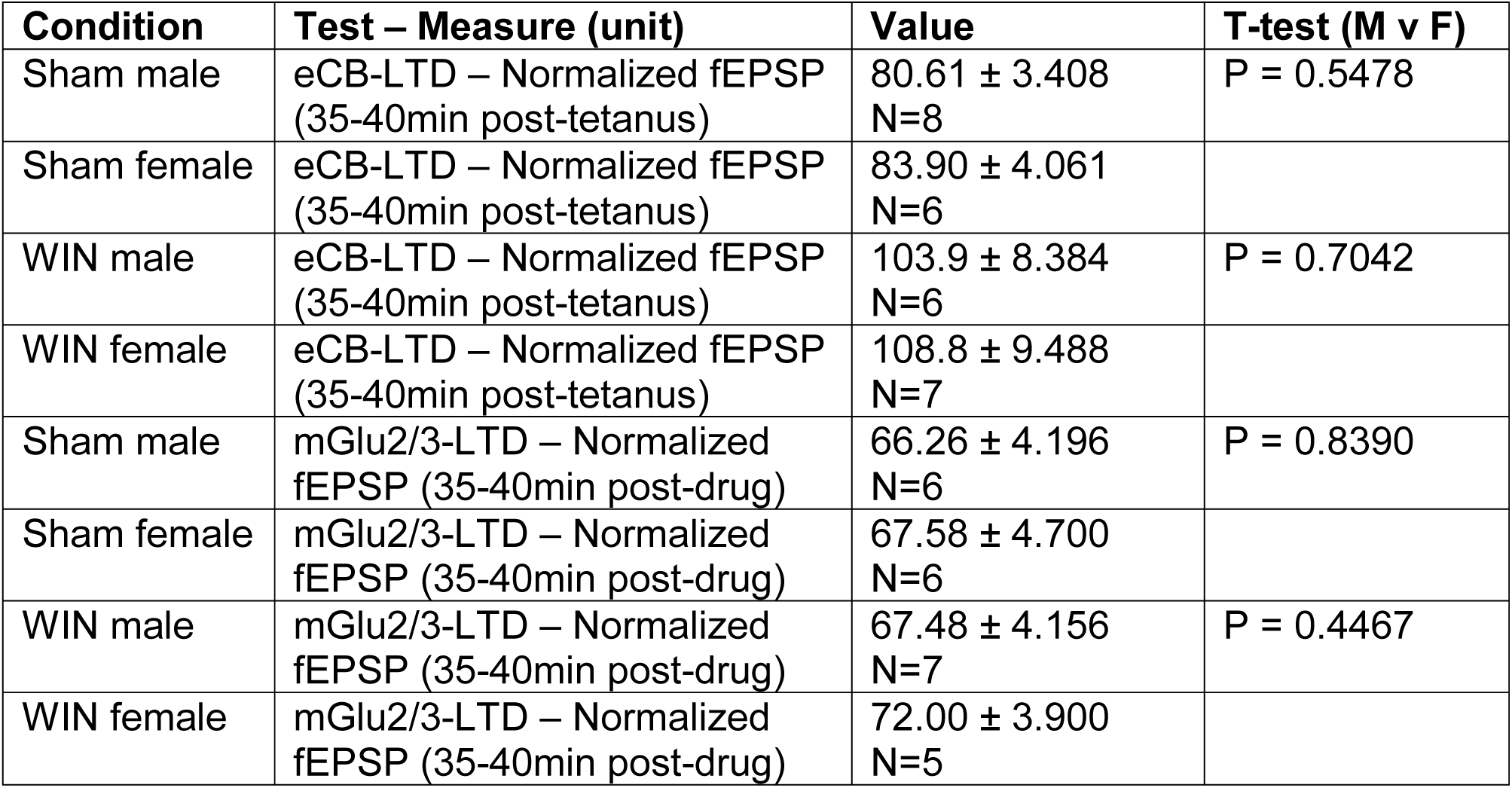
PFC LTD data by sex.

| Condition | Test – Measure (unit) | Value | T-test (M v F) |
| --- | --- | --- | --- |
| Sham male | eCB-LTD – Normalized fEPSP (35-40min post-tetanus) | 80.61 ± 3.408<br>N=8 | P = 0.5478 |
| Sham female | eCB-LTD – Normalized fEPSP (35-40min post-tetanus) | 83.90 ± 4.061<br>N=6 |  |
| WIN male | eCB-LTD – Normalized fEPSP (35-40min post-tetanus) | 103.9 ± 8.384<br>N=6 | P = 0.7042 |
| WIN female | eCB-LTD – Normalized fEPSP (35-40min post-tetanus) | 108.8 ± 9.488<br>N=7 |  |
| Sham male | mGlu2/3-LTD – Normalized fEPSP (35-40min post-drug) | 66.26 ± 4.196<br>N=6 | P = 0.8390 |
| Sham female | mGlu2/3-LTD – Normalized fEPSP (35-40min post-drug) | 67.58 ± 4.700<br>N=6 |  |
| WIN male | mGlu2/3-LTD – Normalized fEPSP (35-40min post-drug) | 67.48 ± 4.156<br>N=7 | P = 0.4467 |
| WIN female | mGlu2/3-LTD – Normalized fEPSP (35-40min post-drug) | 72.00 ± 3.900<br>N=5 |  |

**Table 5.** PFC LTP data by sex.

| Condition | Test – Measure (unit) | Value | T-test (M v F) |
| --- | --- | --- | --- |
| Sham male | TBS-LTP – Normalized fEPSP (35-40min post-tetanus) | 185.6 ± 26.32<br>N=6 | P = 0.3101 |
| Sham female | TBS-LTP – Normalized fEPSP (35-40min post-tetanus) | 155.4 ± 6.180<br>N=7 |  |
| WIN male | TBS-LTP – Normalized fEPSP (35-40min post-tetanus) | 168.0 ± 11.98<br>N=9 | P = 0.3917 |
| WIN female | TBS-LTP – Normalized fEPSP (35-40min post-tetanus) | 152.6 ± 12.58<br>N=6 |  |

**Table 6.** NAc LTD data by sex.

| Condition | Test – Measure (unit) | Value | T-test (M v F) |
| --- | --- | --- | --- |
| Sham male | TBS-LTP – Normalized fEPSP (35-40min post-tetanus) | 82.37 ± 6.937<br>N=5 | P = 0.2893 |
| Sham female | TBS-LTP – Normalized fEPSP (35-40min post-tetanus) | 71.51 ± 6.589<br>N=5 |  |
| WIN male | TBS-LTP – Normalized fEPSP (35-40min post-tetanus) | 110.9 ± 6.257<br>N=5 | P = 0.0714 |
| WIN female | TBS-LTP – Normalized fEPSP (35-40min post-tetanus) | 95.53 ± 3.039<br>N=4 |  |

**Table 7.** PFC intrinsic properties data by sex.

| **Condition** | **Test – Measure (unit)** | **Value** | **T-test (M v F)** |
| --- | --- | --- | --- |
| Sham male | Resting membrane potential (mV) | -67.95 ± 1.408<br>N=6 | P = 0.3580 |
| Sham female | Resting membrane potential (mV) | -66.11 ± 0.9624<br>N=6 |  |
| WIN male | Resting membrane potential (mV) | -64.12 ± 1.103<br>N=7 | P = 0.2000 |
| WIN female | Resting membrane potential (mV) | -66.22 ± 1.077<br>N=6 |  |
| Sham male | Rheobase (pA) | 86.67 ± 18.06<br>N=6 | P = 0.7003 |
| Sham female | Rheobase (pA) | 76.67 ± 17.64<br>N=6 |  |
| WIN male | Rheobase (pA) | 66.19 ± 16.34<br>N=7 | P = 0.5487 |
| WIN female | Rheobase (pA) | 55.00 ± 7.303<br>N=6 |  |

**Table 8.** NAc intrinsic properties data by sex.

| **Condition** | **Test – Measure (unit)** | **Value** | **T-test (M v F)** |
| --- | --- | --- | --- |
| Sham male | Resting membrane potential (mV) | -83.17 ± 1.463<br>N=5 | P = 0.0491 |
| Sham female | Resting membrane potential (mV) | -78.26 ± 1.684<br>N=9 |  |
| WIN male | Resting membrane potential (mV) | -93.95 ± 3.909<br>N=8 | P = 0.5281 |
| WIN female | Resting membrane potential (mV) | -100.4 ± 8.945<br>N=6 |  |
| Sham male | Rheobase (pA) | 199.2 ± 36.64<br>N=5 | P = 0.5827 |
| Sham female | Rheobase (pA) | 176.7 ± 10.93<br>N=9 |  |
| WIN male | Rheobase (pA) | 190.6 ± 20.19<br>N=8 | P = 0.7663 |
| WIN female | Rheobase (pA) | 180.0 ± 28.28<br>N=6 |  |

**Table 9.** Sucrose preference data by sex.

| **Condition** | **Test – Measure (unit)** | **Value** | **T-test (M v F)** |
| --- | --- | --- | --- |
| Sham male | Water consumed (ml) | 1.425 ± 0.1181<br>N=4 | **P = 0.0045** |
| Sham female | Water consumed (ml) | 3.575 ± 0.3326<br>N=4 |  |
| WIN male | Water consumed (ml) | 1.075 ± 0.2287<br>N=4 | P = 0.6953 |
| WIN female | Water consumed (ml) | 1.175 ± 0.04787<br>N=4 |  |
| Sham male | Sucrose consumed (ml) | 3.550 ± 1.533<br>N=4 | P = 0.1069 |
| Sham female | Sucrose consumed (ml) | 7.000 ± 0.7153<br>N=4 |  |
| WIN male | Sucrose consumed (ml) | 8.800 ± 1.564<br>N=4 | P = 0.5118 |
| WIN female | Sucrose consumed (ml) | 6.850 ± 2.291<br>N=4 |  |
| Sham male | Sucrose preference (ratio) | 2.739 ± 1.376<br>N=4 | P = 0.6530 |
| Sham female | Sucrose preference (ratio) | 2.039 ± 0.3597<br>N=4 |  |
| WIN male | Sucrose preference (ratio) | 8.674 ± 1.744<br>N=4 | P = 0.3462 |
| WIN female | Sucrose preference (ratio) | 5.885 ± 2.089<br>N=4 |  |

### Perinatal exposure to a synthetic cannabimimetic alters social behavior and memory at adulthood

In order to determine if the behavioral repertoire of WIN-exposed animals is altered at adulthood, we performed several behavioral analyses in both male and female rats. Based on our previous findings that perinatally THC-exposed animals exhibited augmented social behavior at adulthood, we initiated a social approach and memory assay (Figure 1a-d; Table 1).

**Figure 1.**
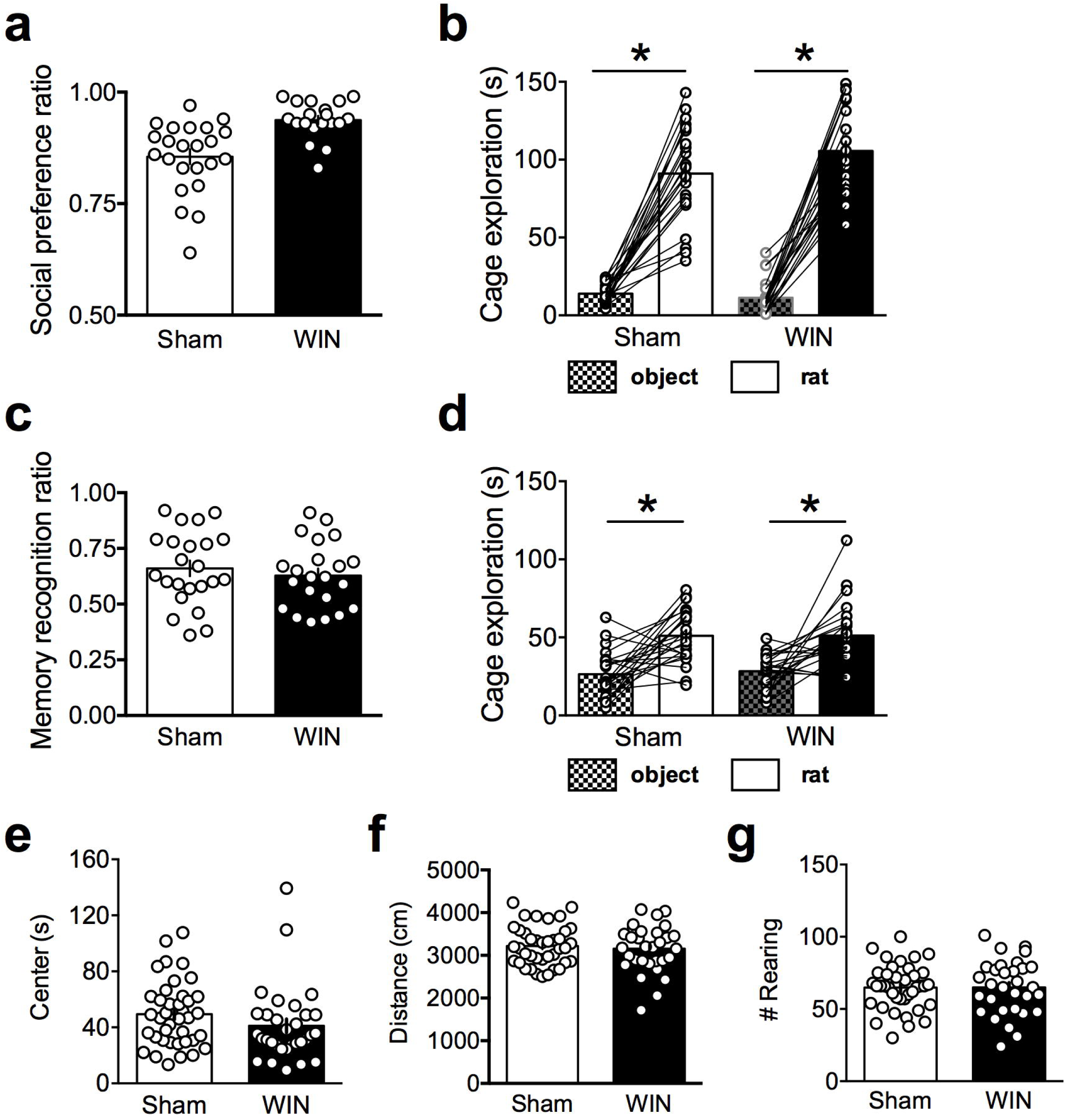
Perinatal WIN alters social approach, but not social memory behavior, nor behavior in the open field environment. **a**: Adult offspring of WIN-treated dams exhibit significantly higher social preference than those of Sham-treated dams (Two-tailed T-test, P = 0.0001; N = 23, 19 respectively). **b**: Time spent exploring a novel rat is significantly higher than time spent exploring a novel object for both Sham- and WIN-exposed rats (One-way ANOVA, F_3,86_ = 14.49; Tukey’s post-hoc analysis, P<0.0001 for both groups; N = 23, 19 respectively). **c,d**: In the subsequent social memory test, the adult offspring of both Sham- and WIN-treated dams exhibited significantly higher preference for a novel, as compared to familiar, rats. **c**: The social memory index does not differ between the two groups (Two-tailed T-test, P = 0.557). **d**: Time spent exploring a novel rat is significantly higher than time spent exploring a familiar rat for both Sham- and WIN-exposed rats (One-way ANOVA, F_3,86_ = 2.137; Tukey’s post-hoc analysis, P<0.0001 for both groups). **e-g**: Behavior in the open field environment does not differ between the offspring of Sham- and WIN-treated dams (N = 39, 31 respectively). Time spent in the center of the arena, total distance covered, and the number of rearing events is not significantly different between groups (Two-tailed T-tests, P = 0.1817, 0.5991 and 0.9783, respectively). *P<0.05

First, during the social approach portion of the assay, WIN-exposed animals exhibited significantly heightened preference for a novel rat over a novel object, as compared to Sham rats (Figure 1a). Both Sham- and WIN-exposed rats did exhibit social preference (Figure 1b), though the magnitude of difference between time spent exploring the novel rat versus the novel object was heightened in the adult offspring of WIN-treated dams. During the subsequent memory test, both Sham- and WIN-exposed rats exhibited a similar social preference for a novel rat over the familiar rat from the social approach assay (Figure 1c-d).

Further, naturalistic behavior was observed prior to the social approach/memory testing by observing animals in the open field assay. No significant differences were noted in the time spent in the center of the arena, distance covered during the trial, or the exploratory behavior (# of rearing events) during the open field test (Figure 1e-g; Table 2).

To determine if other forms of memory were altered by perinatal WIN exposure, we next performed a novel object recognition test. Herein, the adult offspring of both Sham- and WIN-treated dams exhibited similar amounts of object exploration during the initial phase of the test wherein two different, but equally novel objects were presented (Figure 2a; Table 3). However, during the subsequent memory test, while Sham-exposed rats displayed a significant preferential exploration for the novel versus familiar object, WIN-exposed rats failed to discriminate between these two stimuli (Figure 2b; Table 3). Thus, these data indicate that perinatal WIN exposure significantly diminishes novel object memory at adulthood.

**Figure 2.**
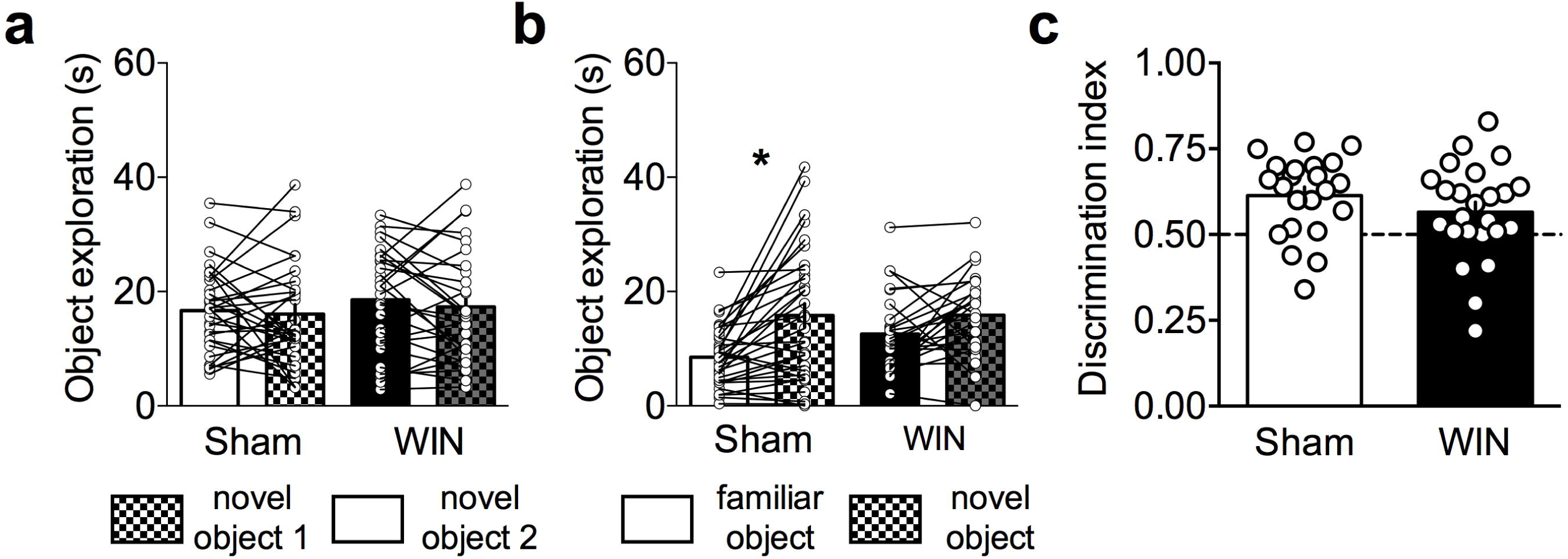
Perinatal WIN exposure abolishes novel object recognition at adulthood. **a**: During training, adult rats from Sham- or WIN-treated dams spent equal amounts of time on two novel objects (One-way ANOVA, F_3,113_ = 0.5446; Tukey’s post-hoc analysis, P = 0.9926 and 0.9485 between objects for Sham and WIN, respectively; N = 30, 28 respectively). **b**: However, during the testing phase, the adult offspring of Sham-treated dams spent significantly more time exploring the novel, over the familiar, object. WIN-exposed offspring did not exhibit significant discrimination between the two objects (One-ay ANOVA, F_3,110_ = 7.970; Tukey’s post-hoc analysis, P = 0.0030 and 0.4867 between objects for Sham and WIN, respectively; N = 30, 28 respectively). **c**: The normalized discrimination index did not differ between Sham- and WIN-exposed offspring (Two-tailed T-test, P = 0.2139).

### Perinatal exposure to WIN alters prefontal synaptic plasticity at adulthood

Previously, we have reported that perinatal THC exposure alters several forms of synaptic plasticity in the PFC at adulthood (Scheyer et al., 2020a). Thus, we elected to examine three forms of PFC plasticity in order to determine if similar alterations followed perinatal WIN exposure. First, we used a 10-minute, 10Hz stimulation of superficial layers of the PFC in order to elicit an endocannabinoid-dependent long-term depression (eCB-LTD) at deep layer synapses (Figure 3a-b; Table 4). Here, we found that while Sham-exposed rats exhibited robust, lasting depression 30-40-minutes following the 10-minute protocol, no such LTD was observed in the PFC of WIN-exposed rats. This finding is in line with our previous data showing an ablation of eCB-LTD in the PFC of THC-exposed rats.

**Figure 3.**
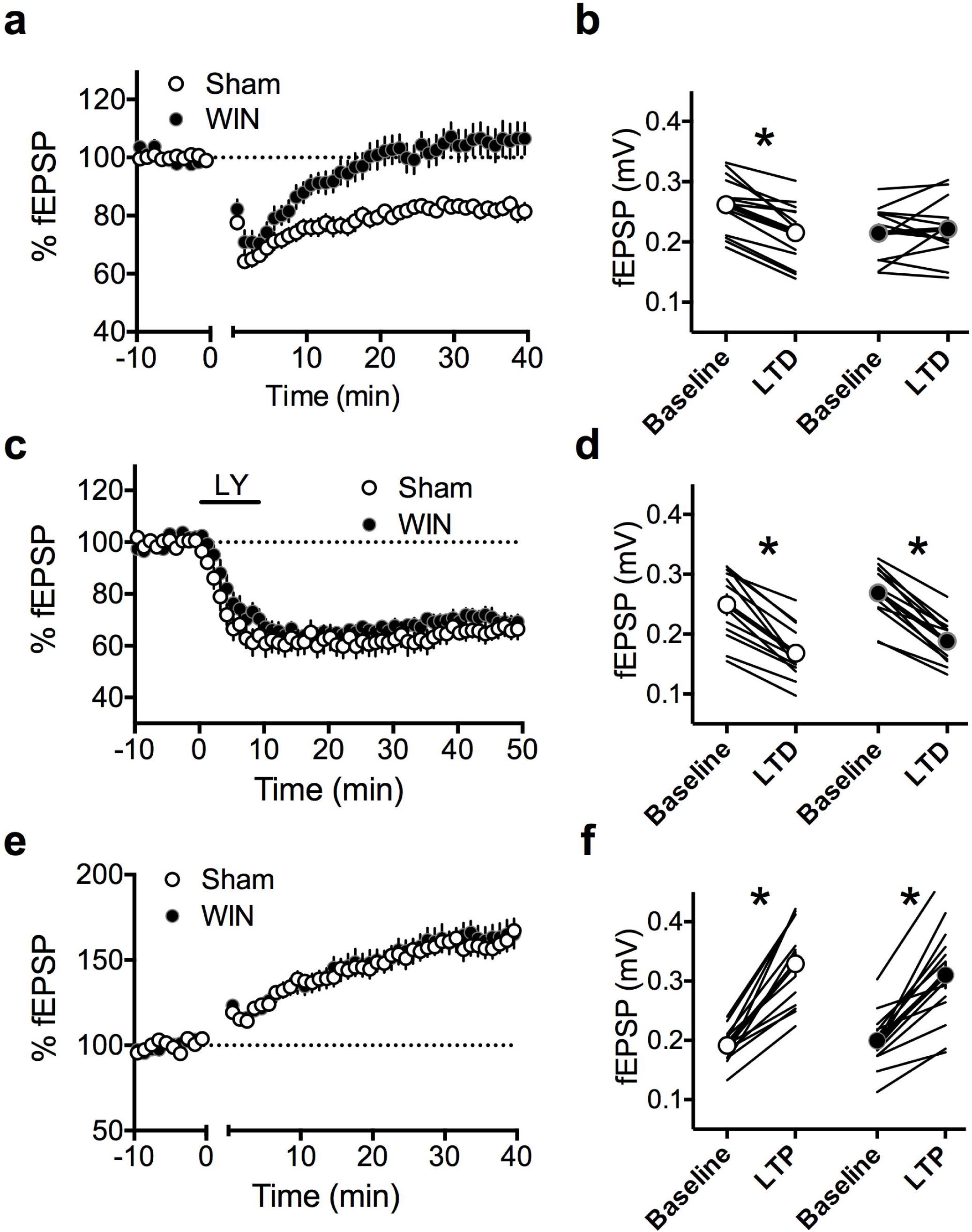
Perinatal WIN exposure induces a selective deficit in LTD in the PFC of adult offspring. **a**: A 10-minute, 10Hz field stimulation of layer 2/3 cells in the PFC of the adult offspring of Sham-treated dams (N = 14) elicited a robust eCB-LTD at deep layer synapses. However, this same protocol failed to induce eCB-LTD in the adult offspring of dams treat with WIN (N = 14). **b**: fEPSP magnitude at baseline (−10 to 0 minutes) and LTD (35-40 minutes post-tetanus) values corresponding to the normalized values in **a** (Two-way RM ANOVA, F_1,24_ = 16.58, P = 0.0004. Sidak’s multiple comparisons test, P=0.0332 and 0.9412, respectively). **c**: LTD mediated by mGlu_2/3_ receptors (mGluR-LTD) is not altered in WIN-exposed offspring. mGluR-LTD, induced via a 10-minute application of LY379268 (LY; 30nM), produced a significant depression at deep layer synapses of the PFC in the adult offspring of both sham- and WIN-treated dams (N = 12 and 12, respectively). **d**: fEPSP magnitude at baseline (−10 to 0 minutes) and LTD (30-40 minutes post-drug) values corresponding to the normalized values in **c**. No differences were found between groups comparing the ten-minute baseline period and the last ten minutes of recording, however both groups exhibited a significant difference of fEPSP magnitude at baseline (i.e. -10 to 0 minutes) as compared to 30-40 minutes post-drug (Two-Way RM ANOVA, F1,11 = 96.69, P<0.0001. Sidak’s multiple comparisons test, P<0.0001 for both groups). **e**: A TBS protocol (5 pulses at 100hz, repeated 4 times) at layer 2/3 cells in the PFC of the adult offspring of both Sham- and WIN-treated dams elicited a robust LTP at deep layer synapses (N = 13, 15 respectively). **f**: fEPSP magnitude at baseline (−10 to 0 minutes) and LTP (30-40 minutes post-TBS) values corresponding to the normalized values in **e**. Both groups exhibited significant differences between the fEPSP magnitude at 30-40 minutes as compared to −10 to 0 minutes (Two-way RM ANOVA, F1,25 = 1.737. Sidak’s multiple comparisons test, P<0.0001 for both groups). *P<0.05

Next, we examined a distinct form of LTD in the PFC mediated by mGlu2/3 receptors (Bara et al., 2018) which has previously been shown to be disrupted by chronic exposure to drugs (Huang et al., 2007; Kasanetz et al., 2013) and augmented at adulthood following perinatal THC exposure (Scheyer et al., 2020a). Thus, we exposed acute PFC slices to the mGlu2/3 agonist LY379268 (300nM) in order to elicit an mGlu2/3-dependent LTD (Figure 3c-d; Table 4). Here, we found that PFC synapses in slices obtained from the offspring of both Sham- and WIN-treated dams exhibited a similar magnitude of mGlu2/3-dependent LTD at 30-40 minutes following drug application.

Finally, we used a theta-burst stimulation protocol at superficial layers of the PFC in order to induce a lasting synaptic potentiation (TBS-LTP) at deep layer synapses. Here, we found no alterations to the time-course or magnitude of plasticity between slices obtained from the adult offspring of Sham-, as compared to WIN-treated dams (Figure 3e-f; Table 5). Of note, these results stand in contrast to those from THC-exposed rats, wherein TBS-LTP is impaired at adulthood (Scheyer et al., 2020a).

Previously, we have found that perinatal THC exposure altered parameters of cell excitability in the PFC at adulthood. Thus, we sought to determine if WIN exposure elicited similar augmentations in excitability. Interestingly, pyramidal neurons in PFC slices obtained from the adult offspring of Sham- and WIN-treated dams did not differ with regards to input-output excitability, spikes elicited by progressive current injections, nor in the rheobase or resting membrane potential (Figure 4a-d; Table 6).

**Figure 4.**
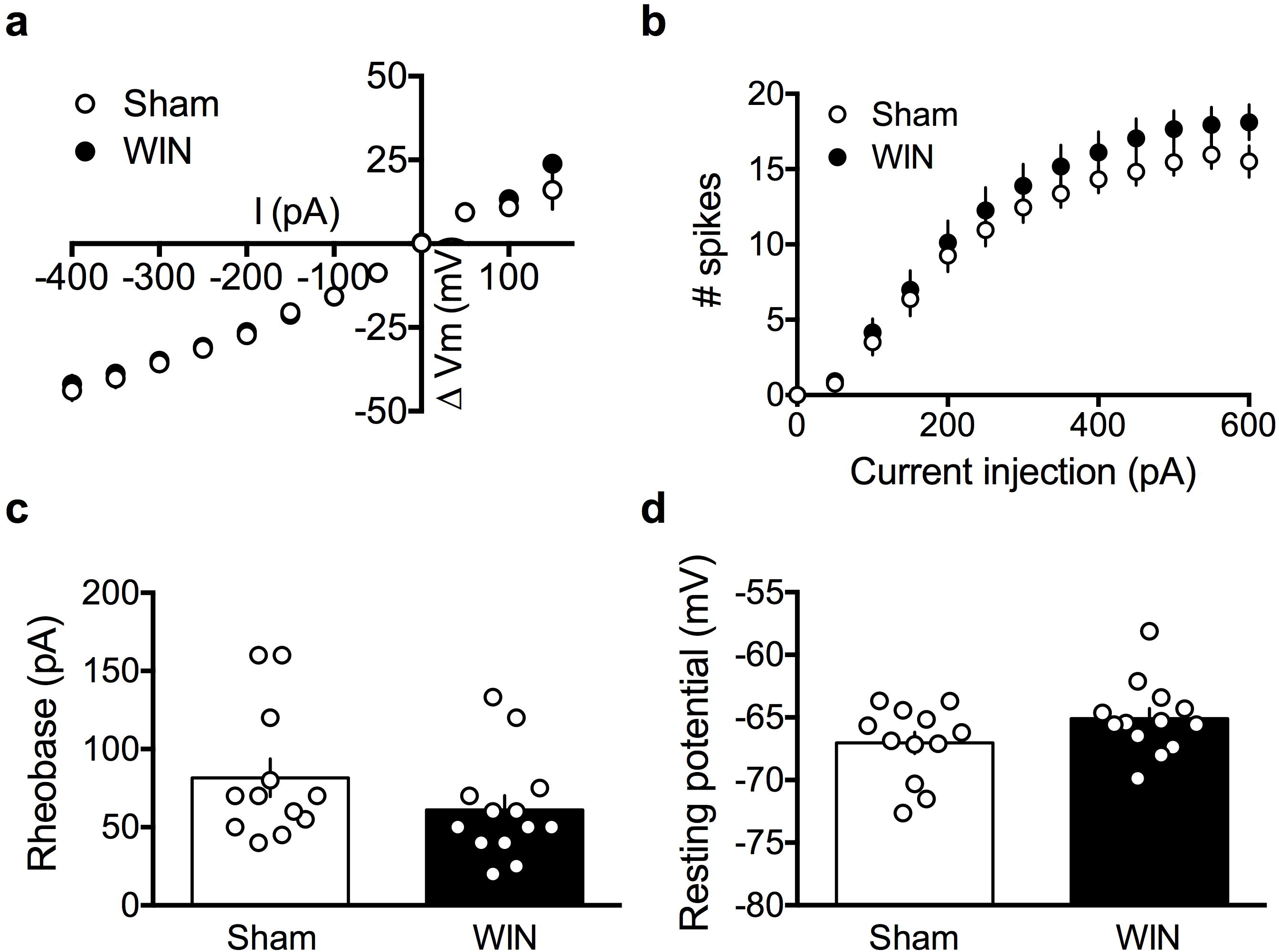
Perinatal WIN exposure does not alter properties of intrinsic excitability of deep layer pyramidal neurons in the PFC of adult offspring. **a**: Current injection steps of 50pA from -400pA to 150pA revealed no differences in the I-V relationship in pyramidal neurons of the PFC between the adult offspring of sham- and WIN-treated dams (N = 12, 13 respectively). **b**: Action potentials elicited by progressive current injections from 0-600pA revealed no difference in the number of spikes elicited in pyramidal neurons of the PFC in slices obtained from the adult offspring of WIN-injected dams as compared to those from sham-treated dams (N = 13, 12 respectively; Two-way RM ANOVA, F_20,460_ = 1.112, P = 0.3328). **c**: Progressive current injections in 10pA steps from 0-200pA revealed that the minimum current injection required to elicit an action potential (i.e. rheobase) did not differ in deep layer pyramidal neurons of PFC slices obtained from the adult offspring of WIN-as compared to sham-treated dams (N = 13, 12 respectively; Two-tailed t-test, P = 0.1896). **d**: Similarly, no difference was found in the resting membrane potential of deep layer pyramidal cells in PFC slices obtained from the adult offspring of WIN-treated dams, as compared to those obtained from sham-treated dams (N = 13, 12 respectively; Two-tailed t-test, P = 0.1123). *P<0.05

### Perinatal exposure to WIN alters synaptic plasticity and cellular properties in the accumbens at adulthood

Recent data have demonstrated that cannabinoids, experimenter- or self-administered, abolish LTD in the NAc (Mato et al., 2004, 2005; Neuhofer and Kalivas, 2018; Spencer et al., 2018). Thus, we sought to determine if perinatal WIN exposure elicited similar deficits in LTD in the NAc at adulthood. We found that while the adult offspring of Sham-treated dams exhibited robust LTD 30-40 minutes after a 10-minute, 10Hz stimulation, no such effect was found in the NAc of WIN-exposed rats (Figure 5a-b; Table 7). Interestingly, unlike in the PFC, these alterations were accompanied by a significant reduction in the resting membrane potential of the principal neurons of the NAc, medium spiny neurons (MSN; Figure 5f). No other parameters of cell excitability were found modified comparing MSNs in slices obtained from Sham-, as compared to WIN-exposed rats (Figure 5c-e; Table 8).

**Figure 5.**
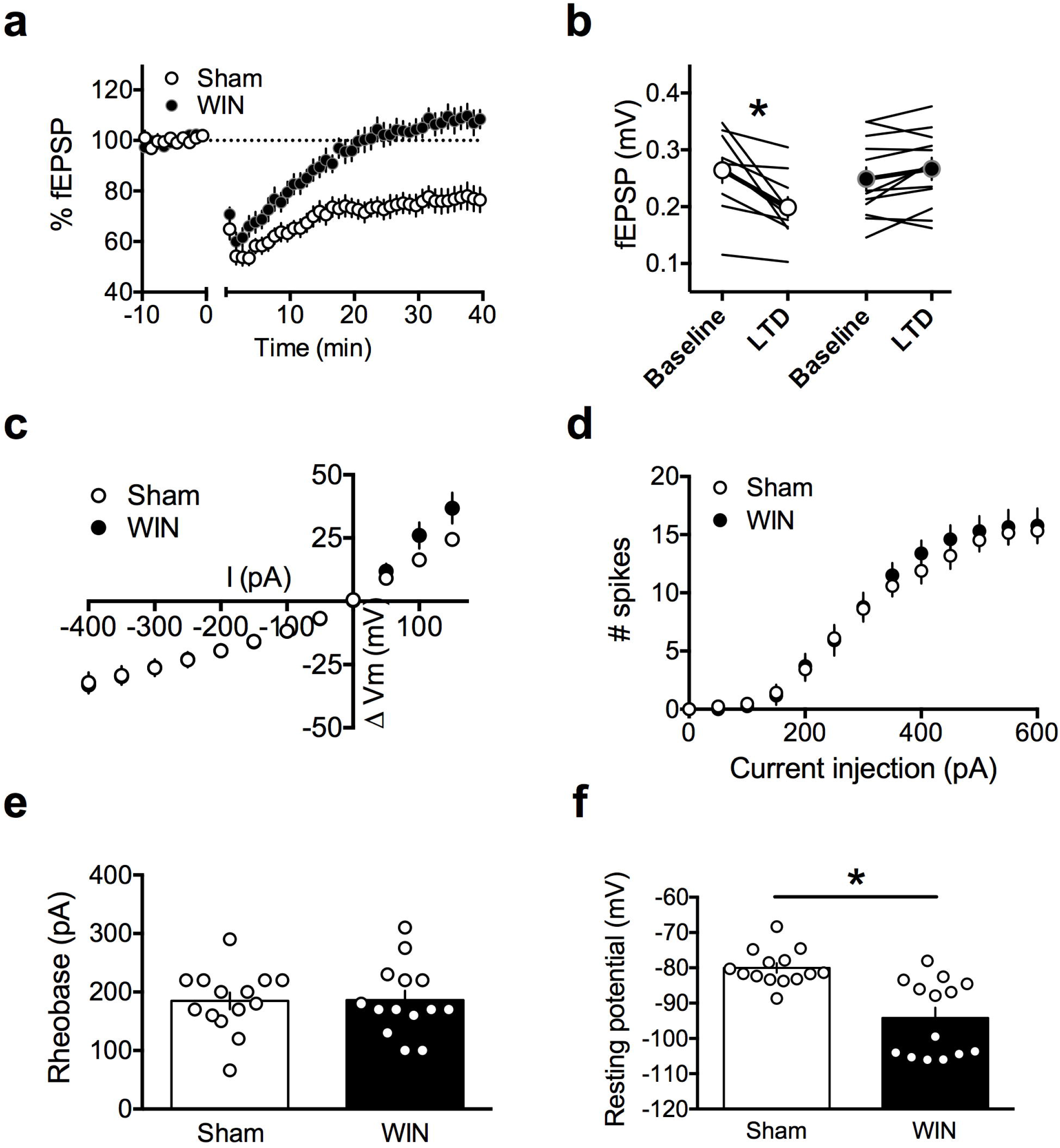
Perinatal WIN exposure abolishes LTD in the NAc of adult offspring and alters the resting membrane potential of NAc medium spiny neurons. **a**: A 10-minute, 10Hz local field stimulation of the NAc of the adult offspring of Sham-treated dams (N = 10) elicited a robust eCB-LTD. However, this same protocol failed to induce eCB-LTD in the adult offspring of dams treat with WIN (N = 12). **b**: fEPSP magnitude at baseline (−10 to 0 minutes) and LTD (35-40 minutes post-tetanus) values corresponding to the normalized values in **a** (Two-way RM ANOVA, F_1,20_ = 20.49, P = 0.0002. Sidak’s multiple comparisons test, P = 0.0002 and 0.3087 for Sham and WIN, respectively). **c**: Current injection steps of 50pA from -400pA to 150pA revealed no differences in the I-V relationship in medium spiny neurons of the NAc between the adult offspring of sham- and WIN-treated dams (N = 14, 14 respectively). **b**: Action potentials elicited by progressive current injections from 0-600pA revealed no difference in the number of spikes elicited in pyramidal neurons of the PFC in slices obtained from the adult offspring of WIN-injected dams as compared to those from sham-treated dams (N = 14, 14 respectively; Two-way RM ANOVA, F_20,250_ = 0.6092, P = 0.9071). **c**: Progressive current injections in 10pA steps from 0-200pA revealed that the minimum current injection required to elicit an action potential (i.e. rheobase) did not differ in medium spiny neurons of NAc slices obtained from the adult offspring of WIN-as compared to sham-treated dams (N=14, 14 respectively; Two-tailed t-test, P = 0.9502). **d**: However, medium spiny neurons in NAc slices obtained from the adult offspring of WIN-treated dams exhibited significantly lower resting membrane potentials than those obtained from Sham-exposed offspring (N=14, 14 respectively; Two-tailed t-test, P = 0.0003). *P<0.05

### Perinatal exposure to WIN enhances sucrose consumption at adulthood

The NAc plays an important role in reward-associated behavior, and recent data indicate a relationship between LTD in the NAc and reward-seeking behavior including sucrose consumption (Bobadilla et al., 2017; Bilbao et al., 2020). Thus, we examined the magnitude of sucrose preference in a two-bottle choice paradigm in the adult offspring of Sham- and WIN-treated dams. Here, we found that while both groups exhibited a preference for a 5% sucrose solution (as compared to plain water) and consumed similar total quantities of liquid during the test (Figure 6a; Table 9), the ratio of sucrose/water consumption was significantly higher in WIN-as compared to Sham-treated adult offspring (Figure 6b; Table 9). Thus, in addition to alterations to synaptic plasticity and intrinsic excitability of MSNs in the NAc, perinatal WIN exposure enhances reward-seeking behavior at adulthood.

**Figure 6.**
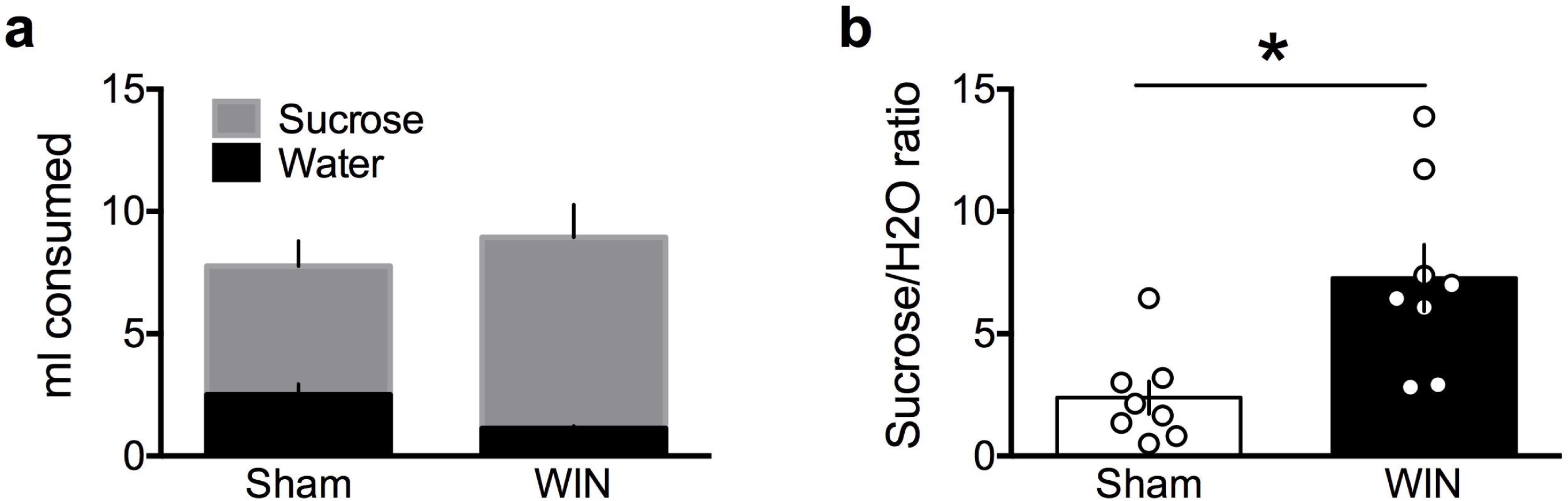
Perinatal WIN exposure increases sucrose preference in adult offspring. **a**: Total quantities of water and 5% sucrose solution (ml) did not differ between the adult offspring of Sham- or WIN-treated dams during a 20-minute sucrose preference test (N = 8, 8 respectively). **b**: Preference for the 5% sucrose solution over water was significantly higher in WIN-as compared to Sham-treated rats (Two-tailed T-test, P = 0.0091).

## Discussion

Here, using the synthetic cannabinoid WIN, we found significant correlations with our previous findings that exposure to plant-derived phytocannabinoid THC via lactation induces behavioral and electrophysiological alterations lasting into adulthood. Specifically, we found altered social behavior, memory, and eCB-mediated synaptic plasticity in the PFC of adult offspring of dams administered WIN during the first 10 days of postnatal life. We also extend the previous data set by showing synaptic deficits and cellular alterations in the NAc along with enhanced sucrose preference, indicative of heightened reward seeking in WIN-exposed adults.

First, our behavioral analyses revealed that perinatal WIN exposure augments social preference in the adult offspring of WIN-treated dams. This result confirms the social augmentation seen following perinatal THC exposure (Scheyer et al., 2020a) but diverge from the effects of in utero THC exposure (i.e. social exploration was reduced, Vargish et al., 2017; Bara et al., 2018). Such discrepancies point to potential differences in the sensitivity of developmental windows through the prenatal and early postnatal periods.

We also report that perinatal despite a lack of differences in social memory, WIN abolishes novel object recognition at adulthood. Interestingly, while object memory is primarily associated with activity in the prefrontal cortex (Barker et al., 2007), social approach and memory is a complex behavior collating activity from diverse brain regions governing motivation and reward such as the amygdala (Adolphs, 2001) and nucleus accumbens (Dölen et al., 2013). Indeed, augmentations in social approach behavior are often associated with decreased amygdalar function and signaling in the nucleus accumbens, where oxytocin-mediated transmission is a key regulator of social approach and reward (Dölen et al., 2013), and is itself governed by the ECS (Wei et al., 2015). Thus, variable impacts on memory and exploration behavior are likely attributable to underlying differences in the driving circuitry.

Results from the current study examining the long-term consequences of perinatal WIN exposure adds to our previous report of dysfunctional eCB-LTD in the PFC of perinatally THC exposed offspring (Scheyer et al., 2020a). In contrast with THC treatment however (Scheyer et al., 2020a), perinatal WIN did not lead to an enhanced magnitude of mGlu2/3-LTD nor a loss of TBS-LTP in the WIN-exposed progeny at adulthood. Differences in the pharmacokinetics, bioavailability and pharmacological profiles of WIN and THC may explain these differences. Despite these subtle differences, these data and those from previous studies suggest that alteration of PFC synaptic plasticity and social behavior at adulthood are common endophenotypes of perinatal cannabinoid exposure (Vargish et al., 2017b; Bara et al., 2018; Scheyer et al., 2019, 2020a).

Previously, we observed that perinatal THC exposure decreases excitability of principle neurons of the PFC (Scheyer et al 2020b) in a fashion similar to chronic adolescent THC exposure in mice (Pickel et al., 2019). Here, we found that no such differences followed perinatal WIN exposure. These data point to a dissociation between measures of intrinsic excitability and synaptic plasticity within the PFC, as changes in these domains appear independent. Thus, alternative explanations for the loss of eCB-LTD must be considered in light of a lack of changes to cell-excitability, including alterations to receptor function or other changes to the ECS such as alterations in eCB tone.

The NAc is essential to reward-associated behavior and we recently showed that eCB-mediated LTD in the NAc core controls reward-seeking behavior (Bilbao et al., 2020). Here, we found that this eCB-LTD is ablated in the NAc of the adult offspring of WIN-treated dams. This finding is in line with multiple reports of altered LTD in the NAc of cannabinoid-exposed animals (Mato et al., 2004, 2005; Neuhofer and Kalivas, 2018; Spencer et al., 2018). In contrast with our recordings in PFC principal neurons, we observed a significant reduction in the resting membrane potential of NAc MSNs. Further, in examining the reward-seeking behavior of these WIN-exposed offspring, we also found that the ratio of sucrose/water consumption in the two-bottle choice task was significantly higher in WIN-as compared to Sham-treated adult offspring. Thus, in the NAc of WIN-exposed progeny, the loss of eCB-LTD and associated cell-excitability modifications were paralleled by modifications of reward-seeking behavior at adulthood.

In conclusion, these results indicate that perinatal exposure via lactation to a synthetic cannabinoid reproduces some of the long-lasting deficits induced at multiple scales by THC. Augmented social behavior and a loss of eCB-LTD in the PFC are therefore similar consequences of perinatal exposure to both naturally occurring phytocannabinoids and synthetic cannabimimetics. Additionally, we found that WIN exposure ablates eCB-LTD in the NAc, where the resting membrane potential of MSNs was found to be significantly decreased. These findings may indeed correlate with an enhanced sucrose-preference amongst WIN-exposed offspring. Together, these findings further illustrate the vulnerability of the developing brain and, consequently, behavior, to early-life insults to the endocannabinoid system via exposure to cannabinoid agonists.

## Author contributions

A.S., M.B., A.L.P and O.J.M. designed research; A.S. and M.B. performed research; A.S. and M.B. analyzed data; A.S. and O.J.M wrote the paper. The authors declare no conflict of interest.

## Acknowledgements

The authors are grateful to the Chavis-Manzoni team members for helpful discussions.

